# A 21□Color Spectral Flow Cytometry Panel for Comprehensive Immunophenotyping of Rat Tissues and a Companion 9□Color T□Cell Functional Assay

**DOI:** 10.64898/2026.10.13.758716

**Authors:** Zhaoyuan Liang, Yang Bai, Liu Yang, Yanfang Li, Zhengyang Guo, Xintong Zhang, Jianling Yang, Lixiang Xue, Yuqing Wang, Jie Zhang

## Abstract

Although rats are crucial translational research models due to their immunological homology and surgical feasibility, the scarcity of conjugated antibodies has hampered high-dimensional immune profiling. To bridge this gap, we created a synergistic two-panel method that uses full-spectral flow cytometry: a 21-color comprehensive immunophenotyping panel and a 9-color intracellular cytokine staining (ICS) panel. The 21-color panel navigates the limited fluorophore pool and maps innate and adaptive lineages at high resolution. While it can be used to profile neutrophils and monocytes in peripheral blood it reaches its full potential in lymphoid organs, allowing for the simultaneous evaluation of tissue-resident macrophages, dendritic cells, T cell differentiation (including Tregs), and APC function. Based on this phenotypic paradigm, the 9-color ICS panel dissects effector T cell activation and Th1/Th2/Th17 cytokine production. Furthermore, we reserved the PE color during panel creation, allowing for the use of antibodies and probes with PE or comparable fluorophores in conjunction with this panel, hence increasing overall flexibility. This combination technique, designed for rat peripheral blood, spleen, and lymph nodes, combines panoramic phenotyping with functional analysis, offering an adaptable foundation for complicated tissue microenvironments in tumor immunology, rheumatology, and autoimmunity research.

## 1 Panel Presentation

### 1.1 Purpose and Appropriate Sample Types

Rats serve as ideal models for translational research due to their high immunological homology with humans and mice. Their larger size significantly lowers the difficulty of complex surgical interventions (e.g., in arthritis models) and facilitates longitudinal tissue harvesting(1). However, the limited availability of commercial antibodies and conjugated fluorochromes has hindered the development of highly multiplexed, stable immunophenotyping panels in rats, restricting the depth of immune profiling.

To bridge this gap, we developed a synergistic two-panel system leveraging full-spectral flow cytometry: a 21-color comprehensive immunophenotyping panel and a complementary 9-color intracellular cytokine staining (ICS) panel (**Table 1**, **2**). Navigating the limited fluorochrome options, the 21-color panel provides high-resolution mapping of major innate and adaptive lineages. Crucially, while adaptable to peripheral blood for profiling neutrophils, circulating monocytes, and lymphocytes, this panel fully unleashes its panoramic potential in lymphoid organs (spleen and lymph nodes). It enables the concurrent evaluation of tissue-resident macrophages, dendritic cells (DCs), and definitive T cell subset differentiation (including regulatory T cells, Tregs), alongside the assessment of antigen-presentation capacity in key antigen-presenting cells (APCs). Building upon this phenotypic framework, the 9-color ICS panel specifically focuses on effector T cell activation and the functional cytokine secretion profiles of Th1, Th2, and Th17 subsets.

**TABLE 1.** Summary table for application of OMIP.

|  |  |
| --- | --- |
| Purpose | Overcoming the Rat Reagent Bottleneck with a 21-Color Comprehensive Immunophenotyping and 9-Color T Cell Functional Spectral Flow Cytometry Panel |
| Technology | Full spectral flow cytometry |
| Application | Research |
| Species | Rat |
| Source of white blood cells | Blood, Spleen and Lymph nodes |
| Cross-references | OMIP 041 |

**TABLE 2.** Reagents used in this OMIP.

| **Panel 1 21-color** |  |  |  |  |
| --- | --- | --- | --- | --- |
| Specificity | Fluorochrome | Clone | Purpose | Dilution titer |
| Viability dye | Zombie-NIR | N/A | Viability | 1:1000 |
| CD45 | BUV805 | OX-1 | Pan hematopoietic marker | 1:80 |
| TCR$\gamma\delta$ | BUV615 | V65 | $\gamma\delta$ T cell marker | 1:40 |
| CD3 | BV605 | IF4 | Pan T cell lineage marker | 1:80 |
| CD4 | RB744 | OX38 | CD4<sup>+</sup> T cell lineage marker | 1:320 |
| CD8 | RY703 | OX-8 | CD8<sup>+</sup> T cell lineage marker | 1:80 |
| CD25 | BV650 | OX39 | T cell activation, Treg lineage marker | 1:160 |
| FoxP3 | PE-Cy5 | FJK-16s | Treg lineage marker | 1:50 |
| CD44 | BB700 | OX-49 | T cell differentiation | 1:160 |
| CD62L | BUV563 | HRL1 | T cell differentiation | 1:320 |
| B220 | BUV496 | HIS24 | B cell lineage marker | 1:40 |
| CD161 | APC | 3.2.3 | NK cell lineage marker | 1:400 |
| CD43 | PE-Cy7 | W3/13 | Marker of monocytes and granulocytes | 1:320 |
| GRA | FITC | HIS48 | Monocytes differentiation | 1:400 |
| CD11b | BUV737 | OX42 | Myeloid lineage marker | 1:320 |
| CD86 | BV786 | 24F | M1 macrophages marker, | 1:160 |
|  |  |  | Co-stimulation marker |  |
| CD163 | RY610 | HIS36 | M2 macrophages marker | 1:40 |
| XCR1 | AF700 | ZET | Marker of classical dendritic cells | 1:200 |
| CD103 | BV421 | OX62 | Marker of type-1 classical dendritic cells | 1:160 |
| CD80 | RB670 | 3H5 | Co-stimulation marker | 1:160 |
| MHC-II | Percp-eflour710 | OX17 | Antigen presenting cells marker, Antigen presenting cells activation | 1:160 |
| **Panel 2 9-color** |  |  |  |  |
| Specificity | Fluorochrome | Clone | Purpose | Dilution titer |
| Viability dye | Zombie-NIR | N/A | Viability | 1:1000 |
| CD3 | BV605 | IF4 | Pan T cell lineage marker | 1:80 |
| CD4 | RB744 | OX38 | CD4<sup>+</sup> T cell lineage marker | 1:320 |
| CD8 | RY703 | OX-8 | CD8<sup>+</sup> T cell lineage marker | 1:80 |
| IFN-γ | AF647 | DB-1 | Th-1 and CTL lineage marker | 1:50 |
| IL-4 | PE | OX81 | Th-2 cells lineage marker | 1:50 |
| IL-17 | FITC | eBio17B7 | Th-17 cells lineage marker | 1:50 |
| CD25 | BV650 | OX39 | T cell activation, Treg lineage marker | 1:160 |
| FoxP3 | PE-Cy5 | FJK-16s | Treg lineage marker | 1:50 |

Together, these panels integrate panoramic immune phenotyping with targeted T cell functional analysis. The protocols have been optimized for rat peripheral immune compartments, including peripheral blood, spleen, and lymph nodes. Given its incorporation of broad phenotypic and functional markers, this combined strategy holds strong potential to serve as a versatile backbone for adaptation to complex tissue microenvironments, particularly in tumor immunology, rheumatology, and autoimmunity research.

### 1.2 Background

The primary objective of this study was to establish a systematic, reproducible, and high-dimensional spectral flow cytometry framework for rats. Given the scarcity of rat-specific reagents, marker selection required balancing cross-species conservation with rat-specific idiosyncrasies to simultaneously achieve panoramic immune profiling and in-depth functional assessment of DC, monocytes/macrophages and T cells (Figure 1A).

**Figure 1.**
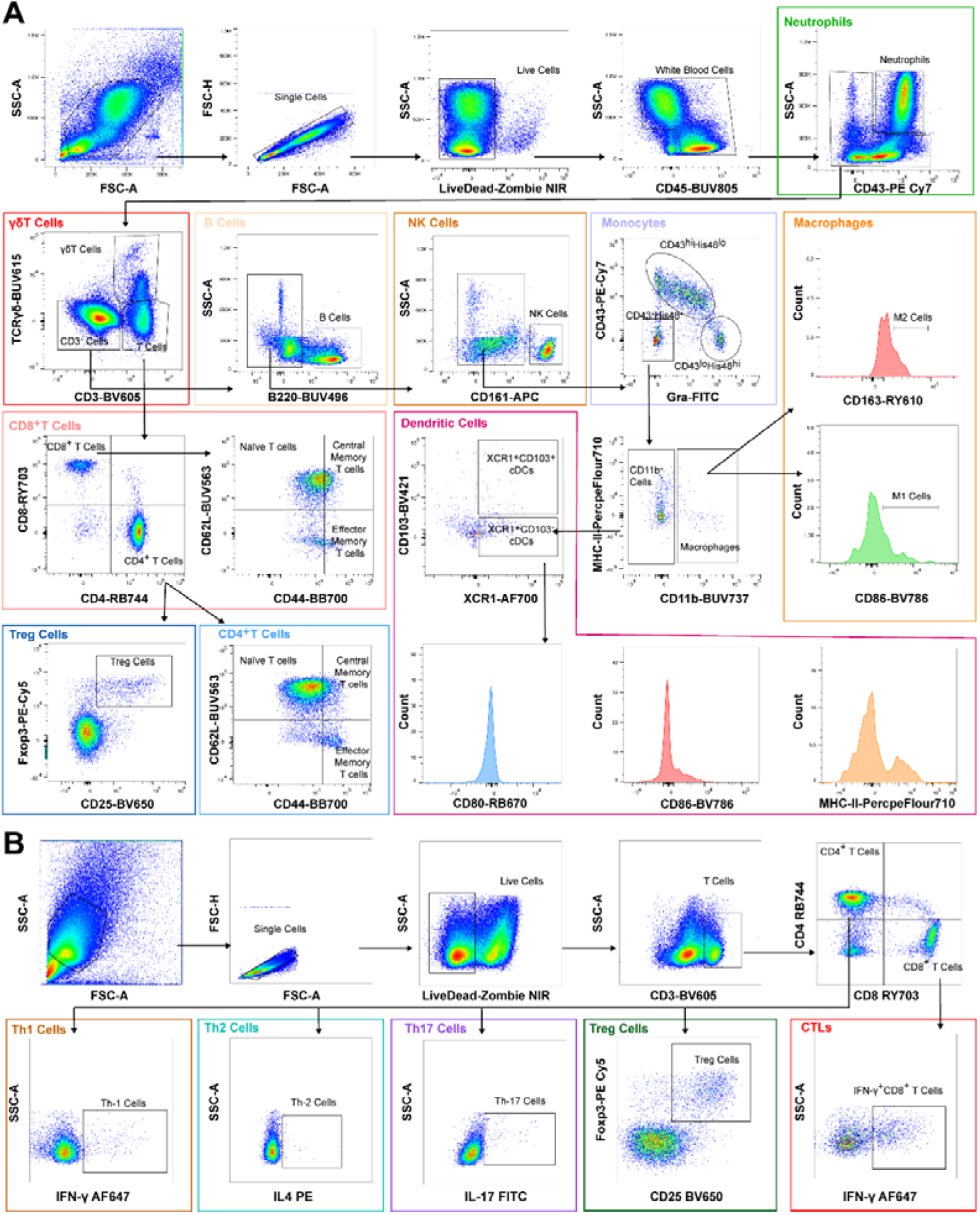
The strategy for the 21-color panel and the 9-color T-cell panel. A: The expression of all markers and the gating strategy for 21-color panel. B: The expression of T cell markers and the gating strategy for 9-color panel.

The myeloid lineage was stratified into granulocytes, monocytes/macrophages, and DCs, utilizing strategies tailored to rat immunobiology. Unlike mice (Ly6G) or humans (CD15), rat neutrophils were definitively identified by their high side scatter (SSC) coupled with CD43 expression(2,3). For monocytes, CD43 and His48 (GRA) expression effectively recapitulated the murine Ly6C paradigm: CD43^Hi^His48^+^ and CD43^Lo^His48^+^ cells were gated as classical and non-classical monocytes, respectively(4,5). Tissue-resident macrophages, identified within the CD43^Lo/-^His48^-^CD11b^+^ compartment, were further evaluated for M1/M2 polarization polarization using CD86 and CD163(6). Dendritic cells, the most potent APCs, were stratified into XCR1^+^CD103^-^ and XCR1^+^CD103^+^ conventional DC (cDC) subsets(7,8). The antigen-presenting maturity and functional status of both DCs and macrophages were subsequently evaluated via MHC-II, CD80, and CD86 expression(9).

Lymphoid gating largely conserved canonical markers, with critical rat-specific adaptations. Unlike human and murine B cells (CD19/CD20), rat B cells were strictly defined by B220 expression(10). Natural killer (NK) cells were characterized by a CD3^-^B220^-^CD161^+^ phenotype(3). Within the T cell compartment, γδ T cells were isolated as CD3^+^TCRγδ^+^, while conventional αβ T cells were subdivided into CD4^+^, CD8^+^, double-positive (DP), and double-negative (DN) populations(7,11,12). Differentiation and trafficking status were mapped using CD62L and CD44 to delineate naïve (Tn), central memory (Tcm), and effector memory (Tem) subsets(11). Regulatory T cells (Tregs) were definitively identified by the co-expression of CD25 and the transcription factor Foxp3(13).

To translate phenotype into function, a dedicated 9-color ICS panel was designed (Figure 1B). This panel quantifies effector cytokines to delineate CD4^+^ T helper polarization (IFN-γ for Th1; IL-4 for Th2; IL-17A for Th17) and CD8^+^ cytotoxic T lymphocyte (CTL) activity (IFN-γ)(14). Crucially, CD25 and Foxp3 were retained in this panel to enable concurrent functional tracking of Tregs. Due to the inherent constraints of the available rat fluorophore repertoire, decoupling the phenotypic and functional markers into two synergistic panels was the optimal strategy to maintain high data resolution and avoid spectral spillover.

Ultimately, despite the severe spectral constraints imposed by the limited rat reagent pool, these two panels successfully close the gap between panoramic immunophenotyping and deep functional profiling. To maximize future adaptability, we strategically reserved a dedicated PE channel within the 21-color panel. This deliberate design choice provides end-users with a flexible ‘plug-and-play’ slot to incorporate additional exploratory markers or metabolic probes (e.g., TMRE, MitoSOX) without disrupting the core spectral backbone.

### 1.3 Similarity to Other Published OMIPS

Herein, we present the first OMIP specifically designed for high-dimensional spectral flow cytometry-based comprehensive immunophenotyping in the rat model. While OMIP- 41established a foundational rat panel, our 21-color panel substantially extends this this foundation by enabling deep profiling of both myeloid lineages (macrophage/DC) and T cell differentiation subsets. Conceptually, it mirrors high-parameter murine/human panoramic panels but is uniquely tailored to circumvent the severe fluorophore constraints inherent to rat immunology (e.g., substituting Ly6G/C with CD43/His48). Furthermore, to address a critical void in functional assessments, we developed a 9-color ICS panel that translates established human T-cell functional OMIPs into a rat-optimized format, facilitating the simultaneous delineation of Th1, Th2, Th17, Treg responses. Collectively, these complementary panels uniquely bridge rat phenotypic mapping with robust function analysis in the rat. Of note, a PE channel was deliberately reserved during the initial panel design to accommodate the subsequent incorporation of PE- or equivalent fluorophore-conjugated antibodies or probes, thereby enabling flexible expansion of the panel for additional parameters.

## 2 Results

Application of the 21-color panel successfully resolved the baseline immune landscapes across rat peripheral blood, spleen, and lymph nodes, demonstrating expected anatomical distributions (Figure 2). Within the myeloid compartment, neutrophils exhibited the highest relative abundance in peripheral blood, were sparsely distributed in the spleen, and were virtually absent in lymph nodes. Circulating monocytes maintained similar frequencies in blood and spleen but remained rare in lymph nodes. Tissue-resident macrophages were predominantly enriched in the spleen, which also displayed a notably elevated proportion of M2-polarized (CD163^+^) macrophages. Dendritic cells (DCs) were highly restricted to secondary lymphoid organs, with the XCR1^+^CD103^+^ DC subset constituting the dominant population in both spleen and lymph nodes.

**Figure 2.**
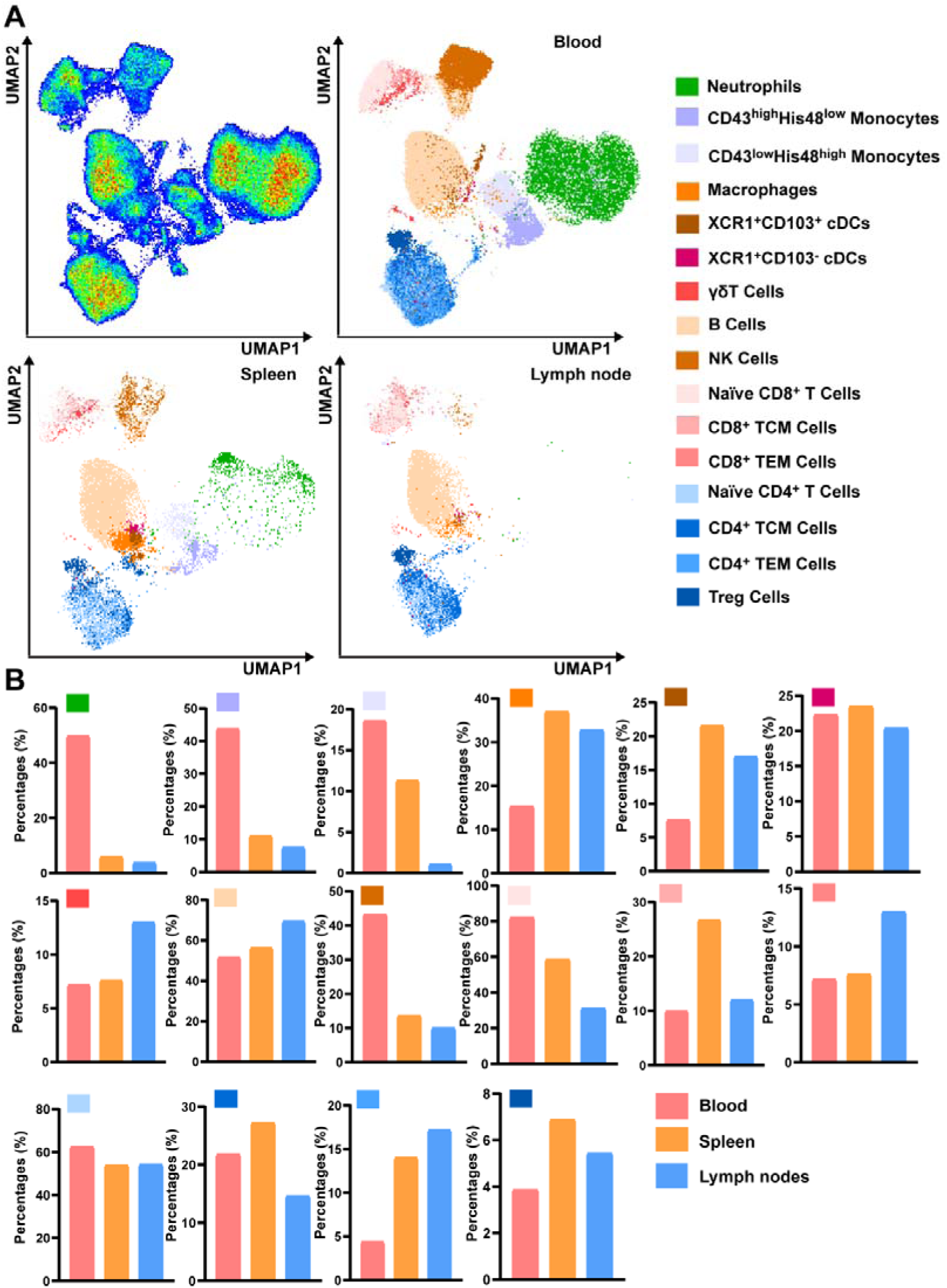
21-color high-dimensional flow cytometry analysis in samples from three sources. A: Visualization of dimensionality reduction results for rat blood, spleen, and lymph nodes. B: Percentages of different immune cells in blood, spleen, and lymph nodes.

In the lymphoid compartment, conventional B and T cells showed a predictable enrichment in the spleen and lymph nodes compared to blood. Conversely, innate-like lymphocytes, the proportion of γδ T cells is higher in lymph nodes, while that of NK cells is highest in the blood. The proportion of Treg cells in the blood is relatively lower than that in the spleen and lymph nodes. These distinct distribution profiles validate the panel’s capability to accurately dissect tissue-specific immune microenvironments.

## 3 Technical Information

### 3.1 Instrument Configuration

#### 3.1.1 Instrument Configuration and Settings

This 21-color flow cytometric panel was developed using a five-laser spectral flow cytometer Sony ID7000 (Sony Biotechnology, Yokohama, JPN). The instrument is equipped with five lasers (355□ nm, 405□ nm, 488□ nm, 561□ nm, and 637□ nm) and 149□(plus FSC and SSC) detectors. As provided in the official Sony ID7000 manual, the instrument scheme is depicted in Figure 3. And the instrument is equipped with 35 UV laser detectors (UV1-UV35), 35 violet laser detectors (V1-V35), 32 blue laser detectors (B1-B32), 26 yellow-green laser detectors (YG1-YG26), and 19 red laser detectors (R1-R19). Detailed detector configuration can be found in Table 5. In principle, the panel can be used on instruments with the same or higher configuration. However, switching to a higher-configuration system (e.g., a 7-laser ID7000) would require testing and possibly adjustments, owing to altered spectral signatures and potential differences in fluorochrome brightness.

**Figure 3.**
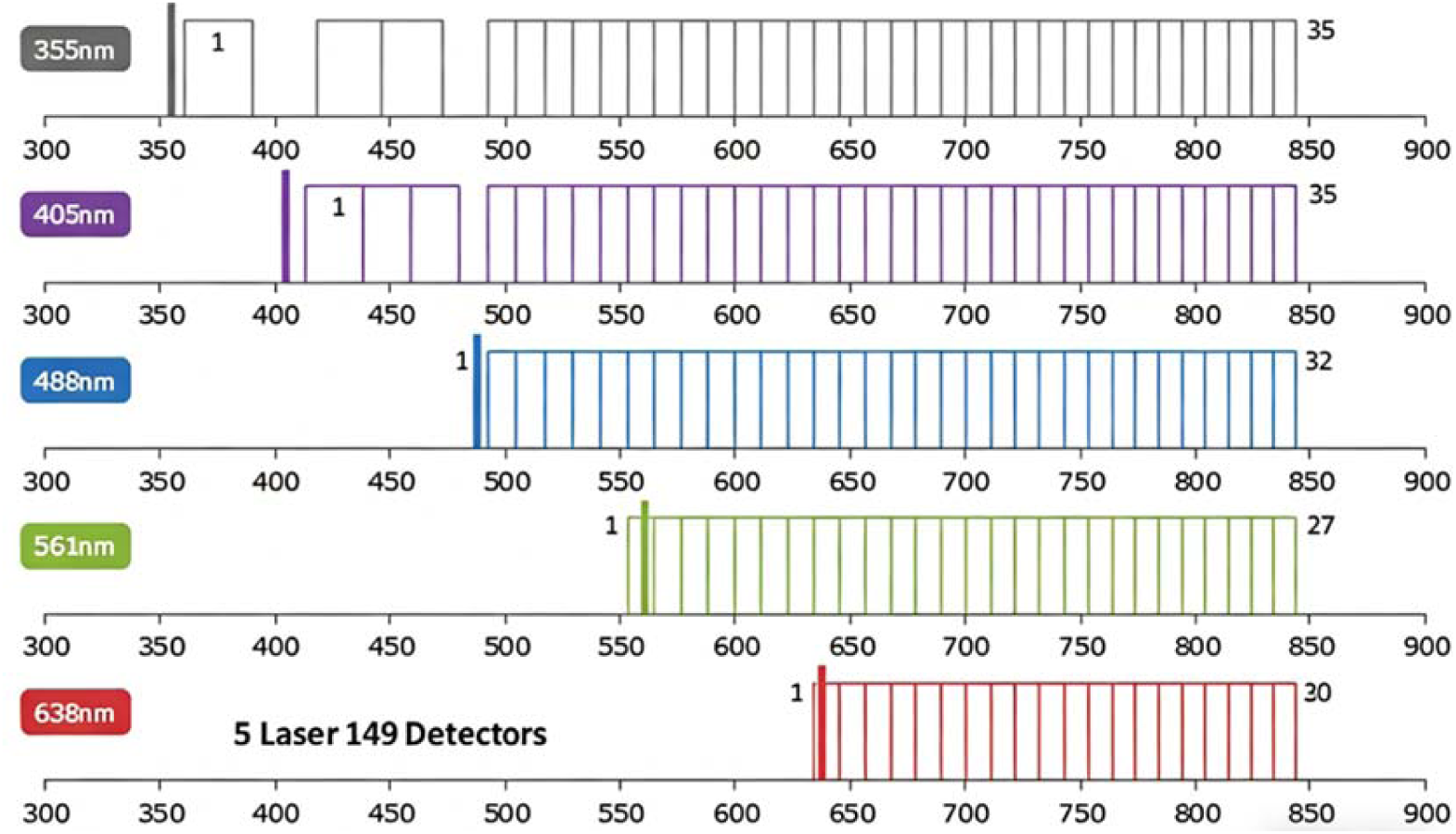
Configuration of ID7000 The SONY ID7000 is equipped with five lasers (355□ nm, 405 □nm, 488 □nm, 561 □nm, and 637□ nm) and 149 detectors, enabling detection of the full fluorochrome emission spectrum.

#### 3.1.2 Quality control

The reliability of measurements from the Sony ID7000 was ensured through a combination of daily QC and performance QC. Commercially available Aligncheck particles were used for daily QC (Sony Biotechnology, JPN, catalog: YAQ02). 8 Peak Beads were used for Performance QC (Sony Biotechnology, Jpn, catalog: YAN01).

#### 3.1.3 Spectral reference generation

Using UltraComp^TM^ compensation control beads (Thermo Fisher, USA, catalog: 01-3333-42), we prepared each fluorophore-conjugated antibody single-stain controls for spectral reference generation. Both beads and flow cytometry antibodies were stained according to the manufacturer’s recommended amounts. The spectral reference of Zombie-NIR was generated by staining cells. After removing autofluorescence and unmixing all spectra, the final flow cytometry data were manually corrected.

### 3.2 Materials

#### 3.2.1 Reagents

The reagents used in this study are listed in Table 3.

**Table 3.** Reagents used for OMIP.

|  |  |  |
| --- | --- | --- |
| Reagent | Cat.No. | Manufacturer |
| PBS (No calcium or magnesium) | SH30256.01 | Cytiva |
| Purified mouse anti rat CD32 | 550270 | BD |
| Horizon Brilliant Stain Buffer (5×) | 568264 | BD |
| FoxP3 Fix/Perm Transcription Factor Staining | 501128857 | eBioscience |
| MACS Buffer | 130-091-221 | Miltenyi |
| Red Blood Cell Lysing Buffer (10×) | 555899 | BD |

#### 3.2.2 Buffers and Prepared Solutions

### Fc Block staining solution: Purified mouse anti rat CD32 antibody was dissolve in PBS. (1:100)

Zombie NIR staining solution (Panel-1 and Panel-2): The powdered Zombie NIR was dissolved in DMSO according to the manufacturer’s instructions, and then diluted 1:1000 in PBS.

Brilliant Stain Buffer: 1□ Brilliant Stain Buffer was diluted from 5× stock with PBS.

Antibody cocktail mix 1 (Panel-1): Dilute the BUV and BV labeled antibodies in 1□ Brilliant Stain Buffer according to the titrated concentration of each antibody.

Antibody cocktail mix 2 (Panel-1): Prepare the other antibodies except BUV- and BV-labeled at appropriate concentrations in PBS.

Fixation/Permeabilization concentrate (1×): 4× stock solution was diluted with Fixation/Permeabilization Diluent. Dilute 1 part concentrate with 3 parts Diluents.

Intracellular stain buffer: 1× Permeabilization Buffer was prepared by the Permeabilization Buffer (10×) diluted in distilled H2O.

Antibody cocktail mix 3 (Panel-1): Foxp3 antibody was diluent in 1× Permeabilization Buffer (1:50 dilution).

Antibody cocktail mix 4 (Panel-2): CD3, CD4, CD8 and CD25 was diluted in PBS and the concentration of antibodies was according to the titrated concentration of each antibody.

Antibody cocktail mix 5 (Panel-2): IFN-γ, IL-4, IL-17 and Foxp3 was diluted in 1×Permeabilization Buffer and the concentration of each antibody was 1:50.

Red Blood Cell Lysing Buffer: 10× RBC Lysing Buffer was diluent to 1× in distilled H2O.

#### 3.2.3 Plastic Consumables

The consumables used in this study are listed in Table 4.

**Table 4.** Plastic Consumables used for OMIP.

| Consumables | Cat.No. | Manufacturer |
| --- | --- | --- |
| 70 µm Cell Strainer | 352350 | Corning |
| 96-well V-bottom plates | E2406336 | Greiner |
| 15 mL centrifuge tube | 430791 | Corning |
| 50 mL centrifuge tube | 430828 | Corning |

**Table 5.**
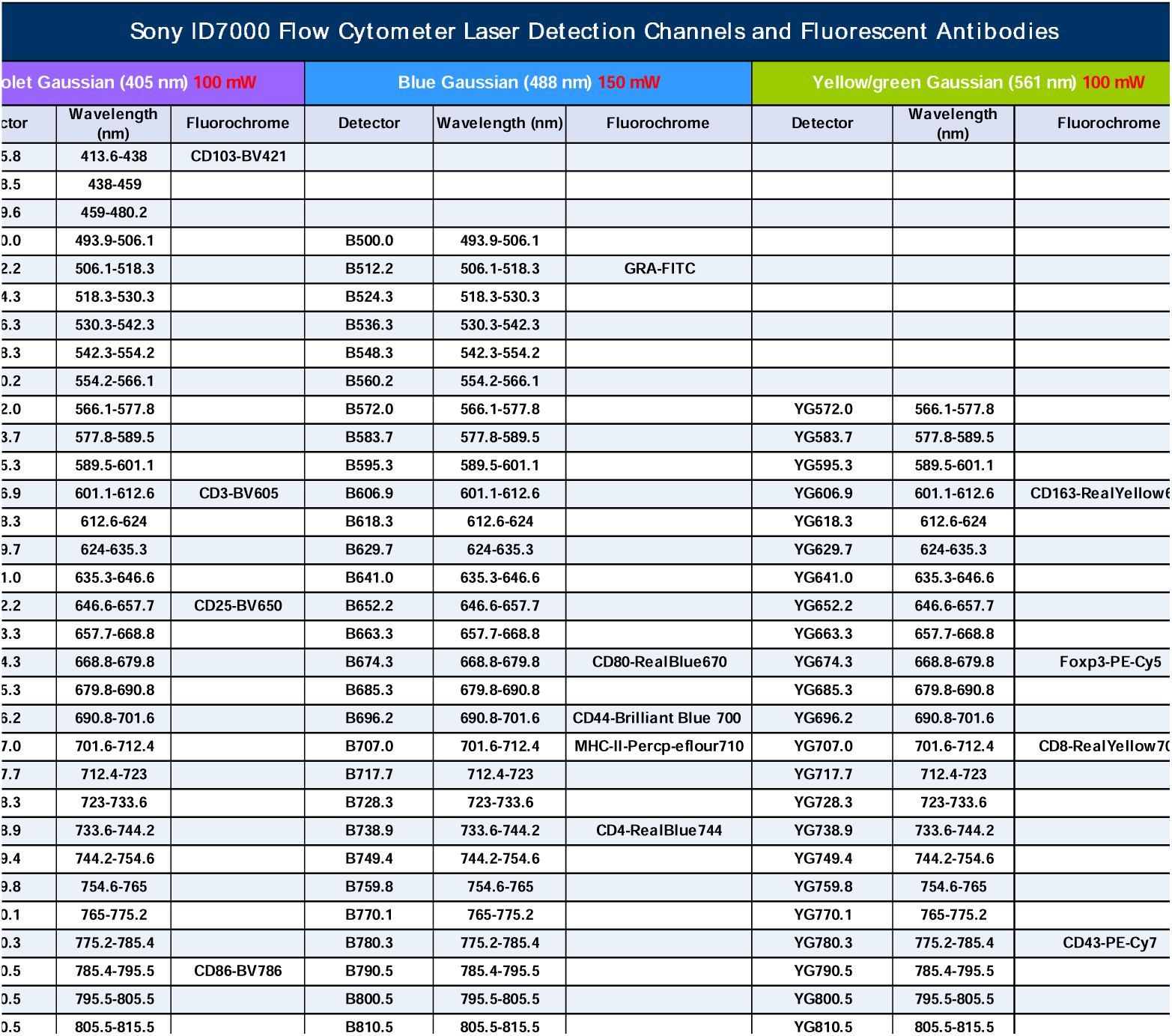
Instrument configuration for the SONY 5 laser ID7000. Sony ID7000 flow cytometer laser detection channels and fluorescent antibodies. The antibodies and fluorophores used in this panel are indicated on the corresponding detectors.

### 3.3 Staining Protocol

#### 3.3.1 Isolation of Single Cells from Rat Spleen, Lymph Nodes, and Peripheral Blood

The study was conducted under the approval of the Animal Ethics Committee of Peking University Third Hospital (Approval No. DLASBE0117). Lymphocytes were harvested from the spleen and lymph nodes as follows: the obtained spleens and lymph nodes were minced, and single-cell suspensions were prepared by grinding through a 70 µm cell strainer. After centrifugation at 500 g for 5 min, the supernatant was discarded, and the pellet was resuspended in 1 × RBC Buffer for 5 min to lyse red blood cells. The lysis was stopped with PBS, followed by centrifugation at 500 g for 5 min. The supernatant was removed, and the cells were resuspended and counted. To obtain leukocytes from peripheral blood, 1 mL of blood which was obtained by sampling from the apex of the heart firstly. And then the blood was diluted with 7 mL of 1× RBC Buffer and incubated for 5 min to lyse red blood cells. The lysis was stopped with PBS, and the mixture was centrifuged at 500 g for 5 min. The supernatant was discarded, and the cells were resuspended and counted. And then, adjust cell concentration to 1×10□ cells per well in 96-well plates.

#### 3.3.2 Full Staining Step-by-Step

##### Panel-1 (21-colors)

1. Cells were plated at 1 × 10□ cells per well in a 96-well plate.
2. Centrifuge samples at 500 g for 5 minutes at room temperature.
3. Remove supernatant and re-suspend the cell in Fc Block staining solution and incubate the cells at room temperature for 30 min.
4. Centrifuge samples at 500 g for 5 minutes at room temperature.
5. Remove supernatant and re-suspend the cell in Antibody cocktail mix1. (BUV and BV labeled antibodies staining)
6. Incubate the cells at 4□ in the dark for 30 minutes.
7. Add 150 µL PBS per well and centrifuge at 500 g for 5 min at room temperature.
8. Remove supernatant and re-suspend the cell in Antibody cocktail mix2. (The other surface markers staining)
9. Incubate the cells at room temperature in the dark for 30 minutes.
10. Add 150 µL PBS per well and centrifuge at 500 g for 5 min at room temperature.
11. Remove supernatant and re-suspend the cell in Zombie NIR staining solution. (Viability Staining Step)
12. Incubate the cells at room temperature in the dark for 15 minutes.
13. Add 150 µL PBS per well and centrifuge at 500 g for 5 min at room temperature.
14. Remove supernatant and re-suspend the cell in Fixation/Permeabilization concentrate (1x)
15. Incubate the cells at room temperature in the dark for 30 minutes. (Fixation/Permeabilization)
16. Add 150 µL 1× Permeabilization Buffer per well and centrifuge at 500 g for 5 min at room temperature.
17. Remove supernatant and re-suspend the cell in Antibody cocktail mix3 for Foxp3 staining.
18. Incubate the cells at room temperature in the dark for 50 minutes. (Intracellular Staining)
19. Add 150 µL 1× Permeabilization Buffer per well and centrifuge at 500 g for 5 min at room temperature.
20. Remove supernatant and Repeat step 17
21. Remove supernatant and re-suspend the cell in 200 μL MACS Buffer. The presence of BSA and EDTA in MACS Buffer ensures that lymphocytes remain as single cells and do not clump, even after extended incubation.
22. Load plate onto the Sony ID7000 and acquire samples.

##### Panel-2 (9-colors)

1. Cells were plated at 1 × 10□ cells per well in a 96-well plate.
2. Centrifuge samples at 500 g for 5 minutes at room temperature.
3. Remove supernatant and re-suspend the cell in Antibody cocktail mix4. (Surface marker staining)
4. Incubate the cells at 4□ in the dark for 30 minutes.
5. Add 150 µL PBS per well and centrifuge at 500 g for 5 min at room temperature.
6. Remove supernatant and re-suspend the cell in Zombie NIR staining solution. (Viability Staining Step)
7. Incubate the cells at room temperature in the dark for 15 minutes.
8. Add 150 µL PBS per well and centrifuge at 500 g for 5 min at room temperature.
9. Remove supernatant and re-suspend the cell in Fixation/Permeabilization concentrate (1x)
10. Incubate the cells at room temperature in the dark for 30 minutes. (Fixation/Permeabilization)
11. Add 150 µL 1× Permeabilization Buffer per well and centrifuge at 500 g for 5 min at room temperature.
12. Remove supernatant and re-suspend the cell in Antibody cocktail mix 5 for intracellular and intranuclear staining.
13. Incubate the cells at room temperature in the dark for 50 minutes. (Intracellular Staining)
14. Add 150 µL 1× Permeabilization Buffer per well and centrifuge at 500 g for 5 min at room temperature.
15. Remove supernatant and Repeat step 17
16. Remove supernatant and re-suspend the cell in 200 μL MACS Buffer.
17. Load plate onto the Sony ID7000 and acquire samples.

#### 3.3.3 Data Acquisition & Software

All flow cytometry data was acquired or analyzed using ID7000 software (Sony Biotechnology). The data acquired on the ID7000 were manually corrected by subtracting autofluorescence and performing manual unmixing correction. And then, the data was and subsequent gating and high dimensional analysis was performed using the flow cytometry software FlowJo™ (BD Biosciences).

## 4 Panel Design and Optimization

### 4.1 Panel Design Strategy

This 21-color panel was designed using a hierarchical, biology-first approach, with a focus on managing the severe reagent restrictions inherent in rat immunology.

To begin, a core backbone of seven markers (CD45, CD3, B220, CD4, CD8, CD43, and MHC-II) was established for global leukocyte classification. Lineage-specific and functional markers were layered to identify myeloid subsets (monocytes, macrophages, DCs) and deep T cell phenotyping (including γδ T cells, Treg identification, and memory differentiation) (Figure 4). Given the limited commercial repertoire of rat-specific antibodies, clone selection was highly influenced by commercial availability and lead times, necessitating preparatory screening to determine optimal clones for this spectral configuration.

**Figure 4.**
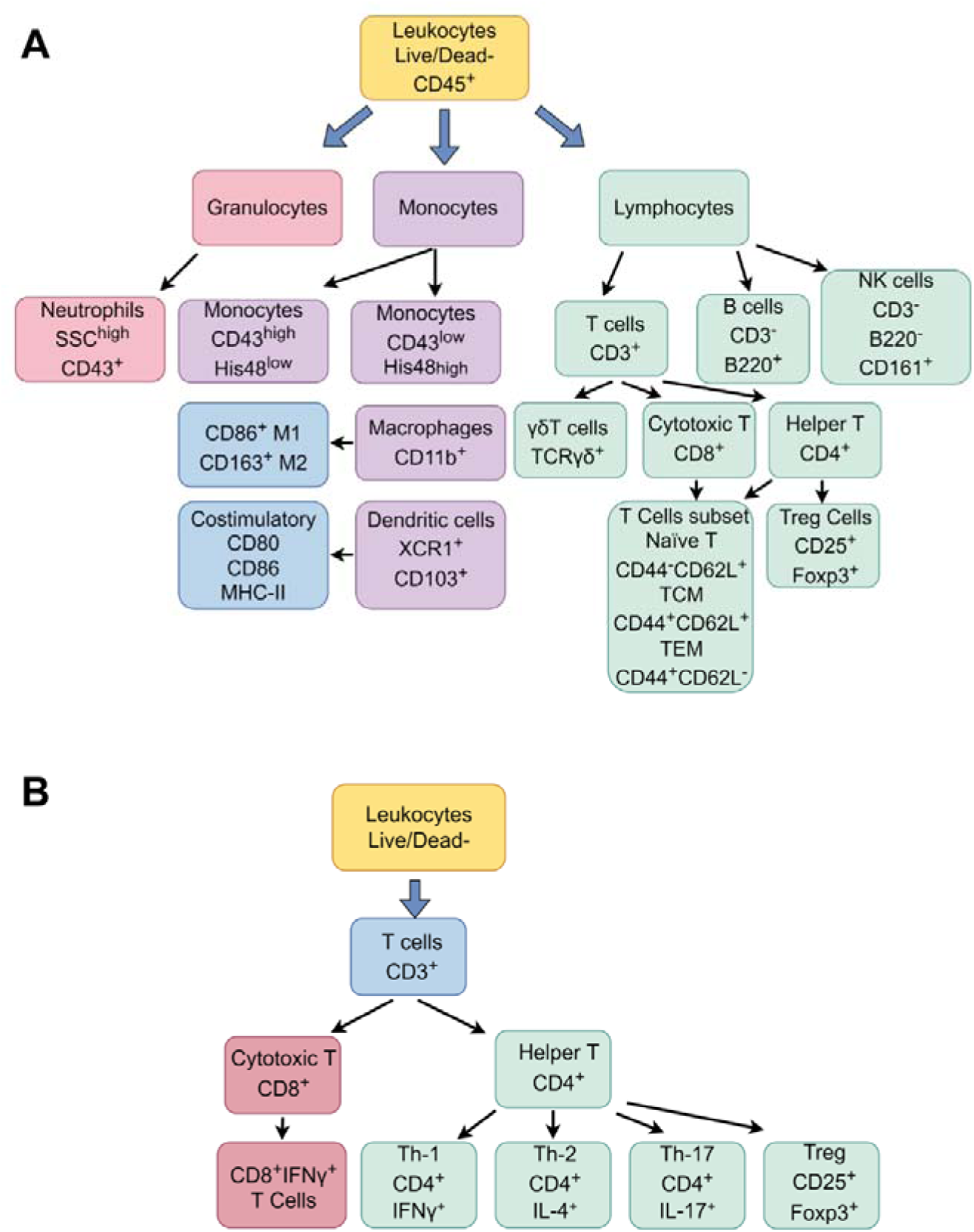
Overview of marker expression on immune cell subsets. A: Expression of markers on different leukocyte subsets (orange): granulocytes (red), monocytes (purple) and lymphocytes (green). Arrows indicate further subset classification. Surface markers on macrophages and dendritic cells are depicted in the blue box. B: Expression of markers on different leukocyte subsets (blue). Arrows indicate further subset classification. Surface markers of CD8^+^ T cells are depicted in the red box. Surface markers of CD4^+^ T cells subsets are depicted in the green box.

Once the biological targets had been determined, strict fluorophore assignment procedures were applied to maximize resolution. The primary rule was antigen-density matching: highly expressed markers (e.g., CD45) were purposely coupled with dim fluorophores (e.g., BUV805), whereas low-density or intracellular targets (e.g., Foxp3) were saved for the brightest available channels (e.g., PE-Cy5). To minimize diffusion spillover, we strictly avoided pairing fluorophores with a high spectral similarity index (SI > 0.85) on co-expressed markers. In contrast, mutually exclusive antigens were strategically assigned to spectrally similar dyes to securely absorb leftover spillover without jeopardizing data integrity.

To overcome the spectral bottleneck caused by the limited rat fluorophore pool, we extensively incorporated innovative BD RealBlue and RealYellow dyes (RB670, RB744, RY703, and RY610). Spectral similarity matrices generated via FluoroFinder directed the minimization of spectral overlap, achieving a low final complexity index of 11.99 (Figure 5). Figure 6 depicts the Spillover Spread Matrix (SSM), which was constructed from single-stained splenocytes at optimal antibody titers and calculated with the FlowJo software version 10.10.

**Figure 5.**
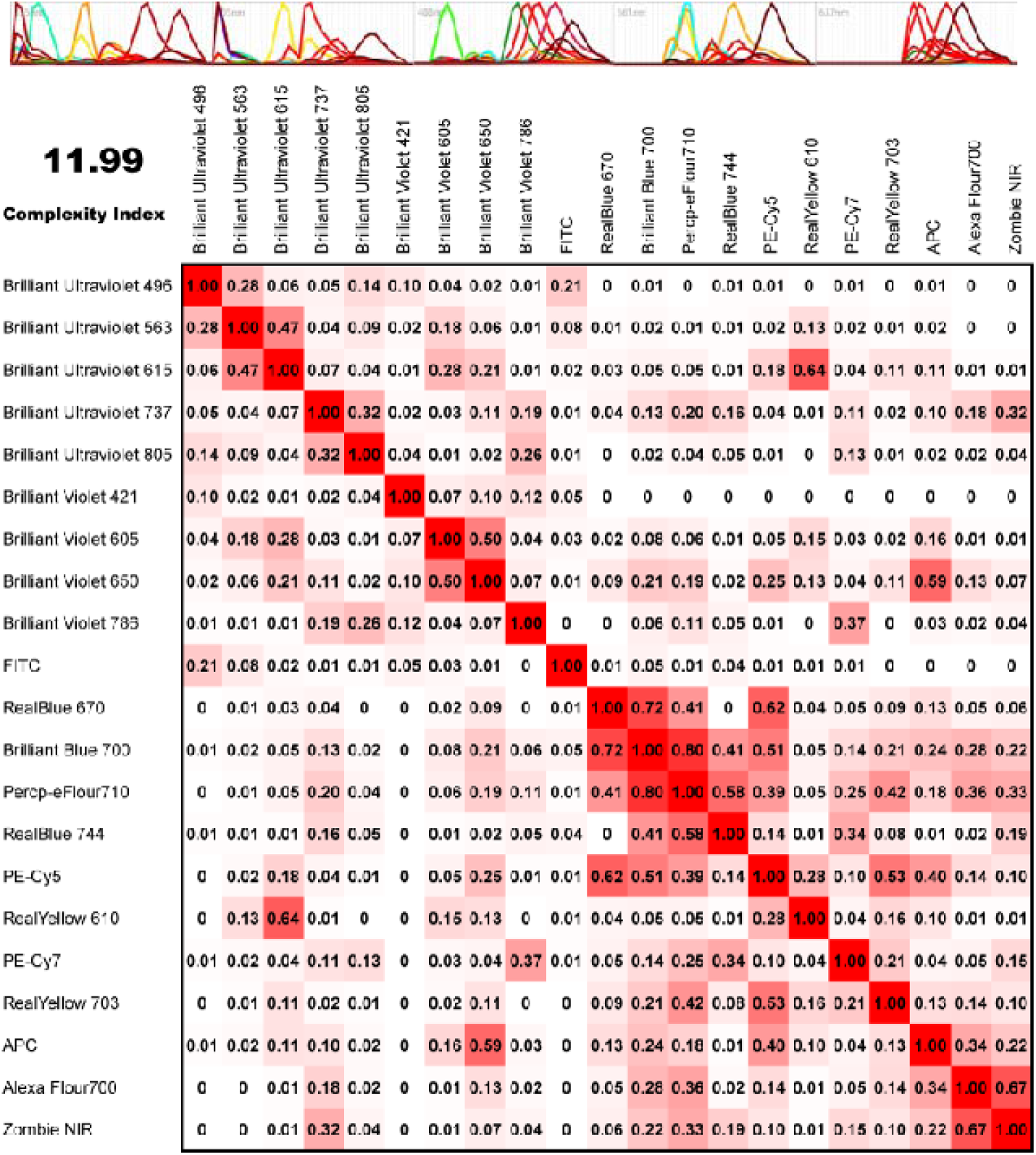
Matrix of similarity indices (SI) for the 21 fluorochromes in the panel. The matrix reports the SI matrix between every pair of fluorochromes in the panel. And the SI matrix was generated using the full spectrum viewer in the FluoroFinder website (https://fluorofinder.com/). The similarity index (SI) is scaled from 0 to 1. A value of 0 denotes no spectral overlap between two fluorochromes, whereas values approaching 1 denote extensive spectral overlap.

**Figure 6.**
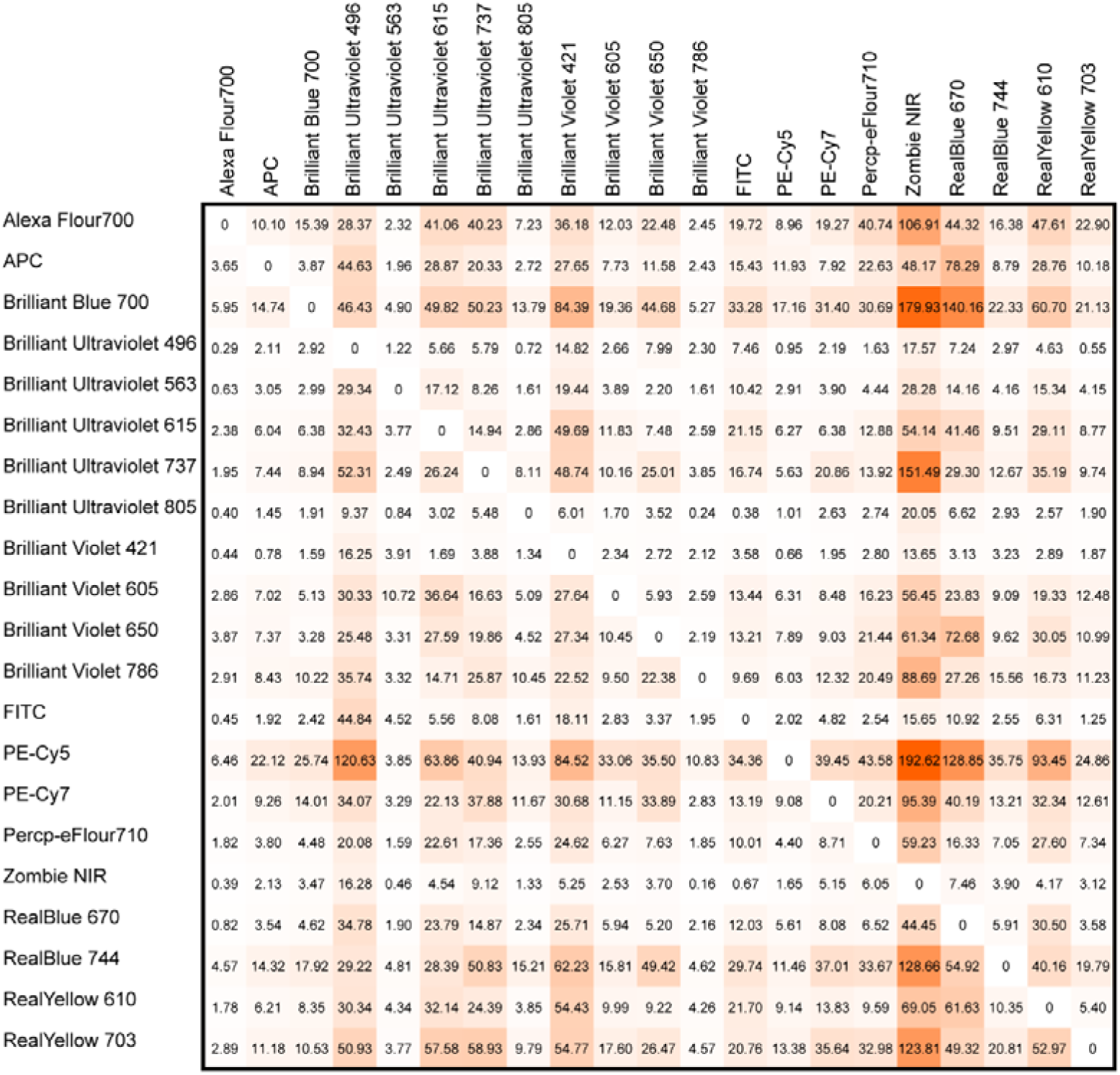
Spreading Spillover Matrix (SSM) in the 21-color panel. The antibodies of spectrums were generated by staining compensation beads individually with 20 different antibody-dye (at the chosen titers). And the spectrum of Zombie NIR was generated by staining the splenocytes. And then the SSM for the final fluorochromes was calculated using Flowjo Software.

Finally, to optimize future adaptability, a dedicated PE channel was purposefully left open. To validate this modular design, we determined spectral similarity indices (using FluoroFinder) for both typical PE-conjugated antibodies and common metabolic probes (e.g., TMRE). The results confirmed that the PE-compatible reagents maintain spectral overlap below the threshold (SI < 0.85) with the existing 21-color architecture (Figure 7), providing a rigorously tested, “plug-and-play” slot for end-users preserving the integrity of the core panel.

**Figure 7.**
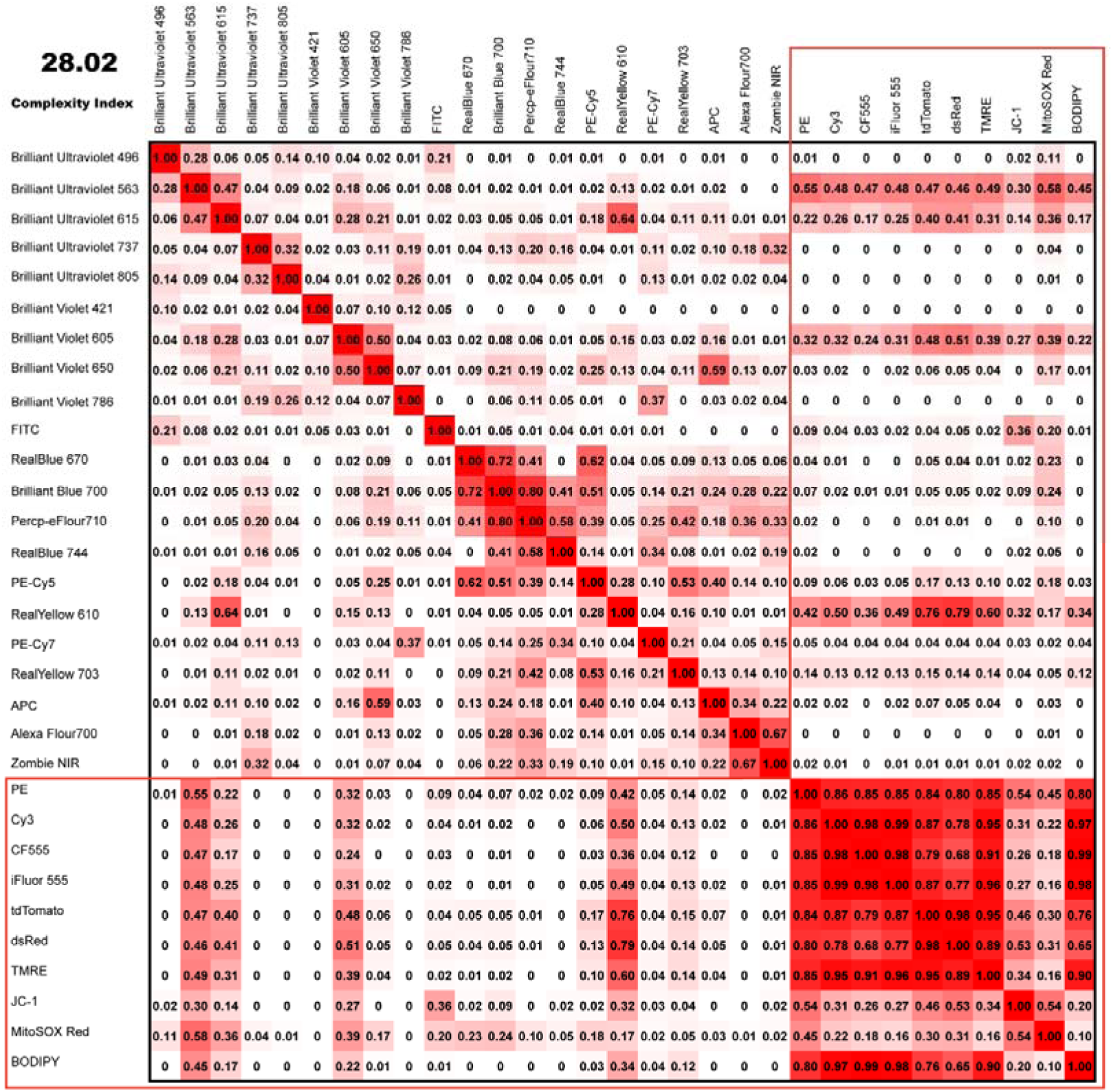
Compatibility of the 21-color panel with PE and similar fluorophores. The red dyes excited by the 561 nm laser like PE, Cy3, CF555, iFlour 555, tdTomato, dsRed, TMRE, JC-1, MitoSOX Red and BODIPY all had similarity values below 0.8 to every fluorophore in the 21-color panel. The probes in the red box are similar to the PE fluorophore and can be freely substituted within this panel.

#### 4.1.1 Selection of Viability Dye

Zombie NIR-LIVE DEAD Zombie NIR dye is excited by a red laser and emits maximally at 746 nm. Zombie NIR was first identified for live/dead discrimination because its far-red properties cause minimal interference with other fluorophores. Notably, Zombie NIR™ dye has an emission spectrum similar to that of APC-Cy7. Consequently, simultaneous use of these two fluorophores should be avoided whenever possible

#### 4.1.2 Fluorochrome Assignment

B220 in BUV496; This dye is moderately to dimly fluorescent and has minimal spread across the panel, making it suitable for a marker like B220, which has a high antigen density.

CD62L in BUV563; This dye has medium-to-high relative brightness and has minimal spread across the panel. Since CD62L is not expressed on all cells and has a moderate expression level, we paired BUV563 with it.

TCRγδ in BUV615; BUV615 is a bright tandem dye well suited for markers with medium-to-low antigen density, such as TCRγδ. Although BUV615 may have some spectral similarity with CD163 in RY610; TCRγδ and CD163 both have low expression and are not co-expressed, which making BUV615 is a good choice for this panel.

CD11b in BUV737; This dye has medium-to-high relative brightness and has minimal spread across the panel. Because its emission spectrum has low similarity to those of other fluorochromes, we matched it with CD11b, which is expressed at a relatively low level.

CD45 in BUV805; Given its low spillover and dim emission, this dye is ideally assigned to markers with high antigen density that are extensively co-expressed, such as CD45.

CD103 in BV421; We assigned CD103 to BV421 due to the low antigen density of CD103 and moderate-to-high brightness of BV421. In addition, BV421 causes minimal spread across the panel, which is ideal for CD103 given its only express in DCs.

CD3 in BV605; BV605 is a moderately bright dye that shows low spillover spreading across the panel. We assigned CD3 to BV605 due to CD3 is a panel backbone marker and is highly expressed.

CD25 in CD650; BV650 is a moderately bright dye that exhibits some spillover into other channels. We assigned CD25 relatively low antigen density to BV650 to minimize the spreading effects.

CD86 in BV786; BV786 is a medium□brightness dye. Because it has low spillover into other fluorochromes and is well suited for antigens with unknown expression levels, we paired CD86 with BV786.

GRA in FITC; FITC derivatives are the most widely used fluorescent reagents in biological research, owing to their high absorptivity and excellent fluorescence quantum yield. We assigned GRA to FITC due to commercially available antibodies against the rat GRA come in almost exclusively this one fluorochrome. If other fluorochromes are available, FITC can be used for markers with lower expression.

CD80 in RB670; RB670 is a novel bright dye with high brightness and low spectral overlap. Which significantly reducing spreading errors as compared to PE-Cy5. Therefore, we paired it with CD80 as CD80 is a marker with relatively low expression.

CD44 in BB700; BB700 is a bright dye well suited for markers with medium-to-low antigen density, such as CD44. Although it has some spectral overlap with Percp□eFluor710 and RB670, we overcame this limitation by pairing it with non□co□expressed markers.

MHC-□ in Percp-eFluor 710; Percp-eFluor 710 has medium-to-high relative brightness. We assigned MHC-□ to Percp-eFluor 710 due to MHC-□ is expressed only on antigen presenting cells and is not co□expressed with CD44.

CD4 in RB744; RB744 is a bright dye that shows low spillover spreading across the panel. We assigned CD4 to RB744 due to CD4 is a panel backbone marker and is highly expressed.

FoxP3 in PE-Cy5; This dye has high relative brightness and has minimal spread across the panel. Since Foxp3 is only expressed on Treg cells and has a dim expression level, we paired PE-Cy5 with it.

CD163 in RY610; RY610 is a bright dye that shows low spillover spreading across the panel. We assigned CD163 to RY610 due to CD163 has a dim expression level in macrophages. Moreover, CD163 is not co□expressed with TCRγδ-BUV615.

CD43 in PE-Cy7; PE-Cy7 is a bright dye that shows low spillover spreading across the panel. And it was assigned to CD43 due to the limited availability of commercial anti-rat CD43 antibodies.

CD8 in RY703; RY703 is a bright dye that shows low spillover spreading across the panel. We assigned CD8 to RY703 due to CD8 is a panel backbone marker and is highly expressed.

CD161 in APC; APC is a bright dye that allows sensitive discrimination of antigens with medium to low expression. Although APC does exhibit spreading interactions with BV650 dyes, we attempted to manage through allocation of CD161 which is a marker only express in NK cells.

XCR1 in AF700; Although Alexa Fluor 700 dye is a dim, near-infrared dye, commercially available antibodies against rat XCR1 are limited.

### 4.2 Panel Optimization

#### 4.2.1 Antibody Titration

All antibodies underwent 2-fold serial dilution titration on fresh rat splenocytes, starting at 2× manufacturer recommended concentration. Cells were plated at 1 × 10□ cells per well in a 96-well plate. And then we added 100 µL of the antibody mix at the appropriate concentration, incubate in the dark for 20 minutes. After 20 minutes, the cells were washed twice with PBS, and then the samples were ready for analysis. Optimal dilutions marked with red boxes in Figure 8, selected for maximal separation of positive/negative populations with minimal background. The final dilution titer of each antibody is shown in Table 2 and Figure 8.

**Figure 8.**
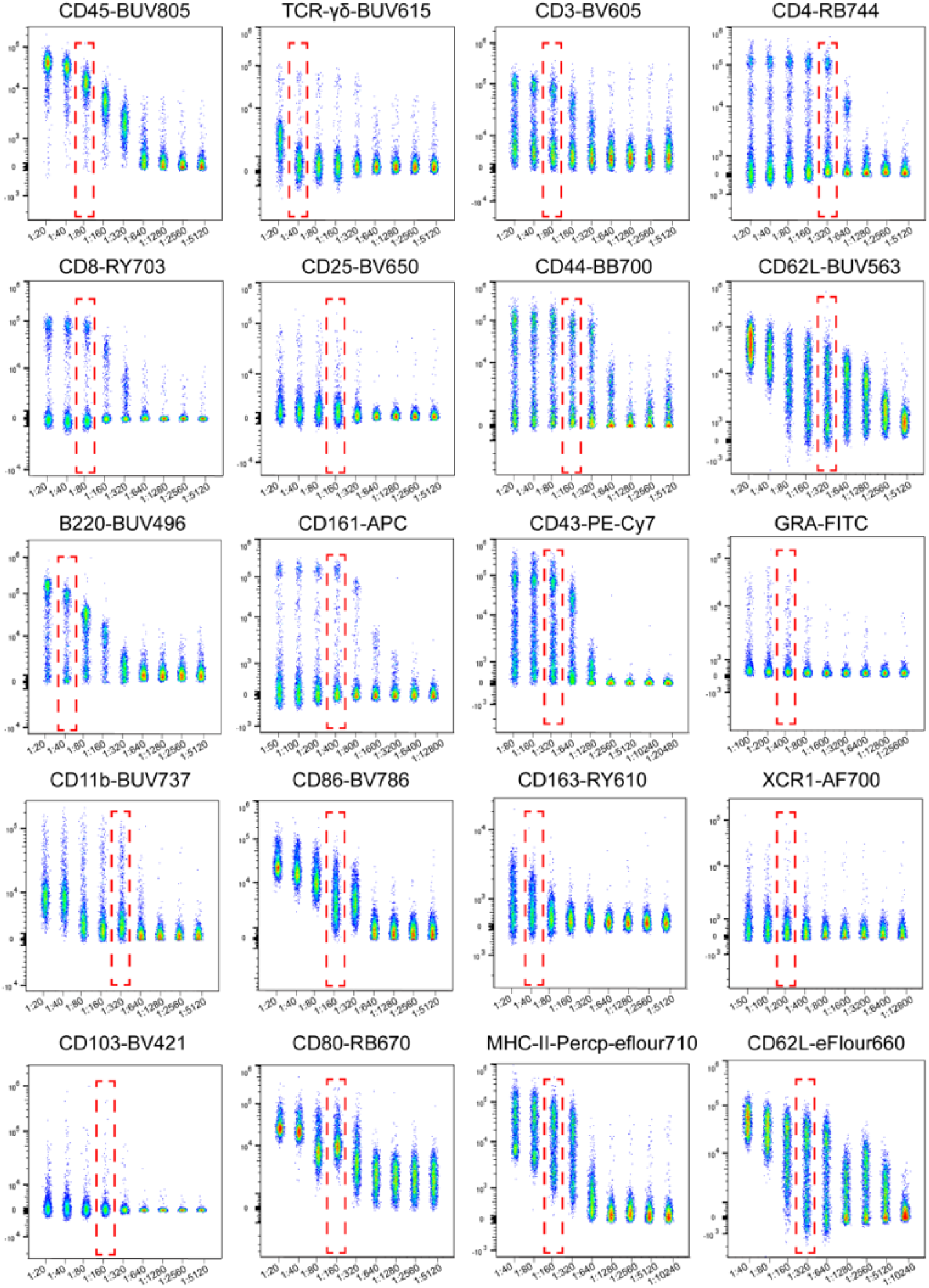
Titration of all antibodies included in the OMIP. Antibodies were titrated in three-fold dilutions on freshly isolated splenocytes of rat. Red rectangles highlight the selected dilution factor used in the OMIP.

#### 4.2.2 Fluorophore Clone Comparison (CD62L example)

CD62L-eFluor660 (Clone: OX85) showed poor population resolution compared to CD62L-BUV563 (Clone: HRL1) in both blood and lymph node samples; BUV563 clone was selected for final panel (Figure 9).

**Figure 9.**
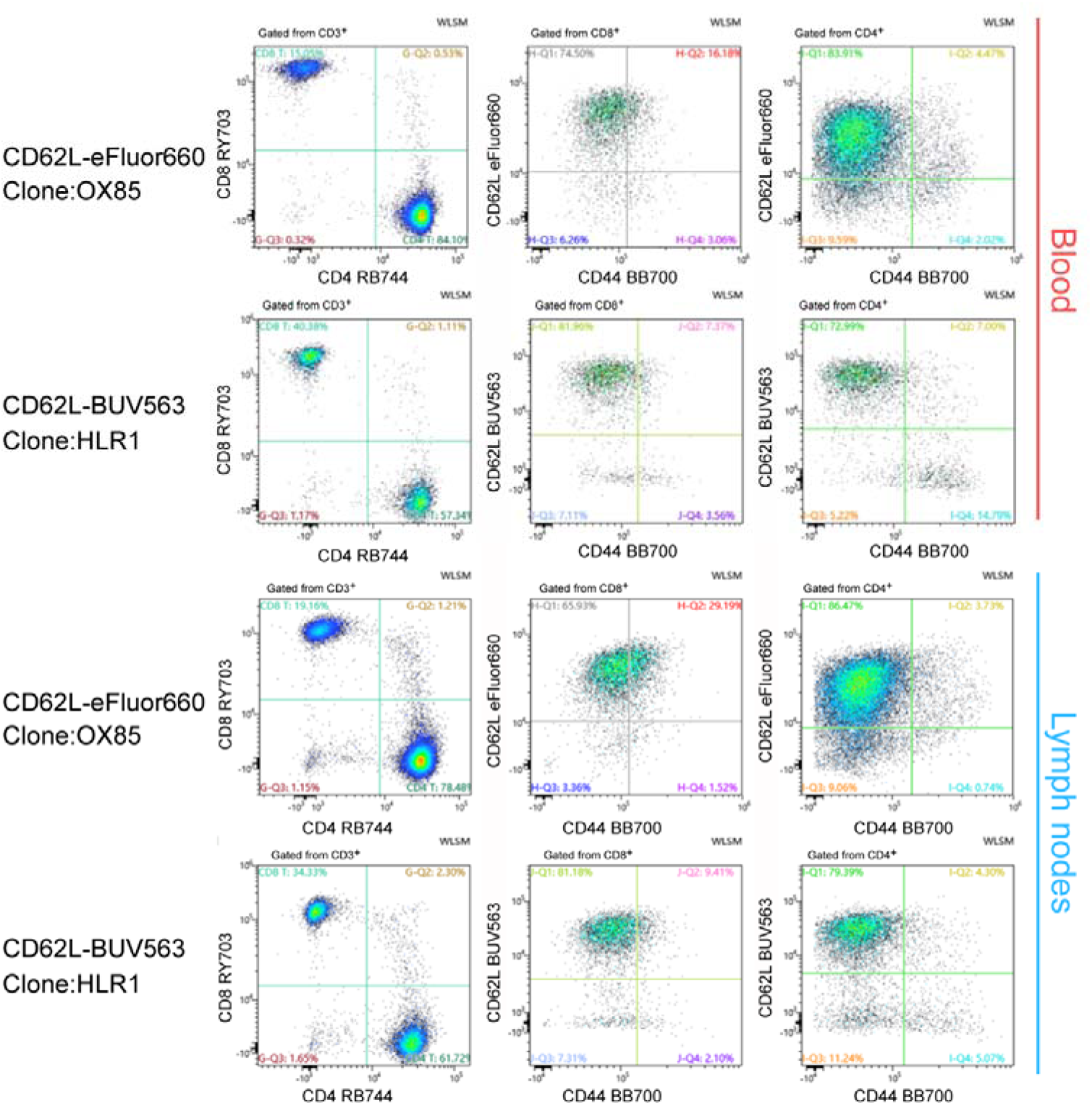
CD62L antibody optimization. In blood (red labeling) and lymph node (blue labeling) staining, we observed that CD62L-BUV563 gave clearer cell population separation than CD62L-eFluor660. We therefore performed antibody optimization and selected CD62L-BUV563 in the OMIP.

### 4.3 Spectral Quality Control

After data acquisition, direct analysis using the established spectra may introduce certain errors, and autofluorescence is a critical factor need to be considered. We observed that autofluorescence excited at 355□ nm and 488□ nm caused more pronounced interference with other fluorophores. Manual quality control of the experimental results was included and autofluorescence was subtracted from the data (Figure 10). In addition, directly unmixed data may also contain errors, and we checked the unmixing results for each fluorophore, using manual N×N plot adjustment to eliminate false populations (Figure 11). To ensure the accuracy of flow cytometry staining, we performed single staining for the antibodies in this panel and compared the single stains with the full panel stain. Histogram overlay confirms minimal loss of resolution after full panel staining (Figure 12).

**Figure 10.**
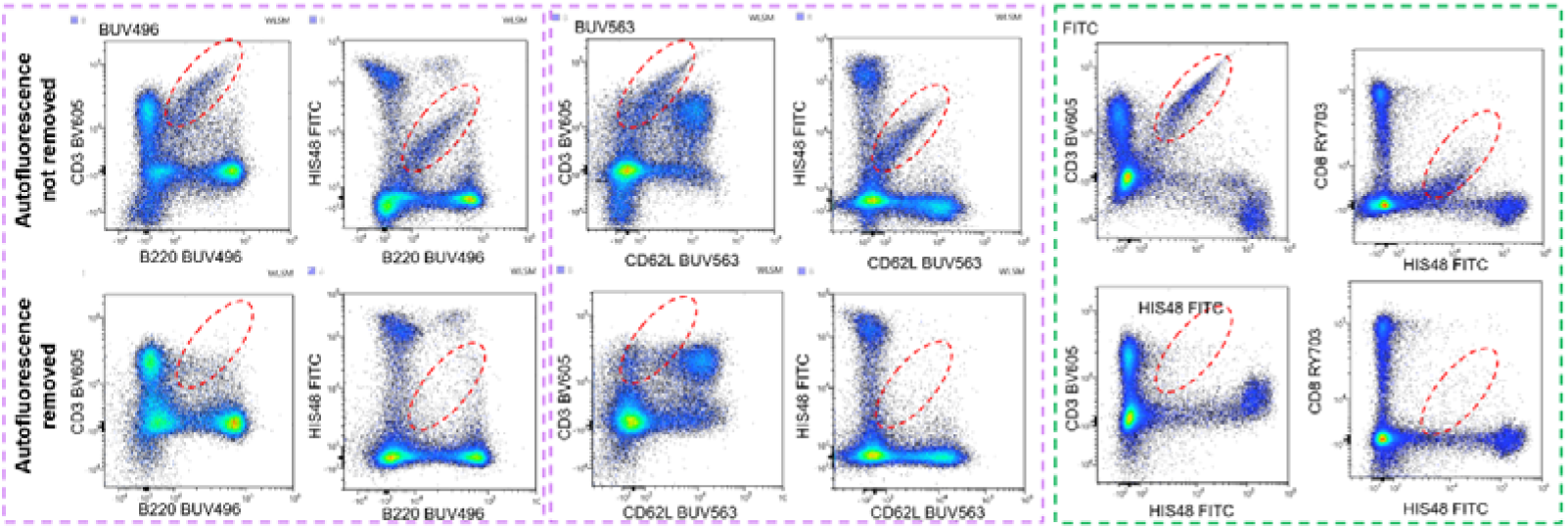
Autofluorescence correction. Autofluorescence subtraction: The fluorescence of BUV496, BUV563 and FITC were depicted as an example on the x-axis. Red boxes marked false positive signals that appeared before autofluorescence subtraction.

**Figure 11.**
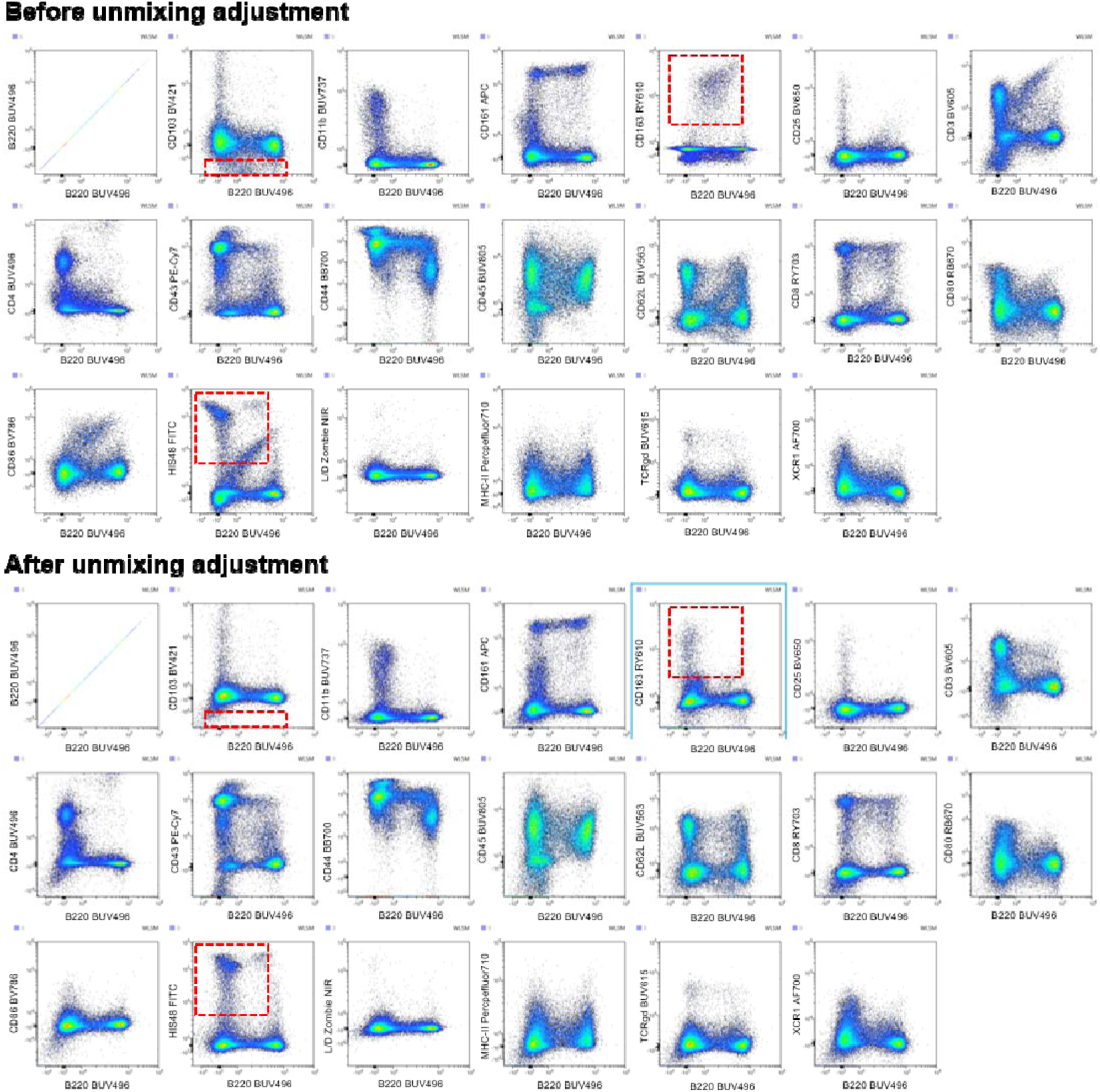
Unmixing and Adjustments: N×N plots viewed with the ID7000™ Unmixing Viewer. Unmixing was performed on the live cell population. Unmixed data with B220-BUV496 depicted as an example on the x-axis and all other fluorochromes plotted on the y-axis. Examples of unmixing errors and corrected unmixing results are indicated by red boxes

**Figure 12.**
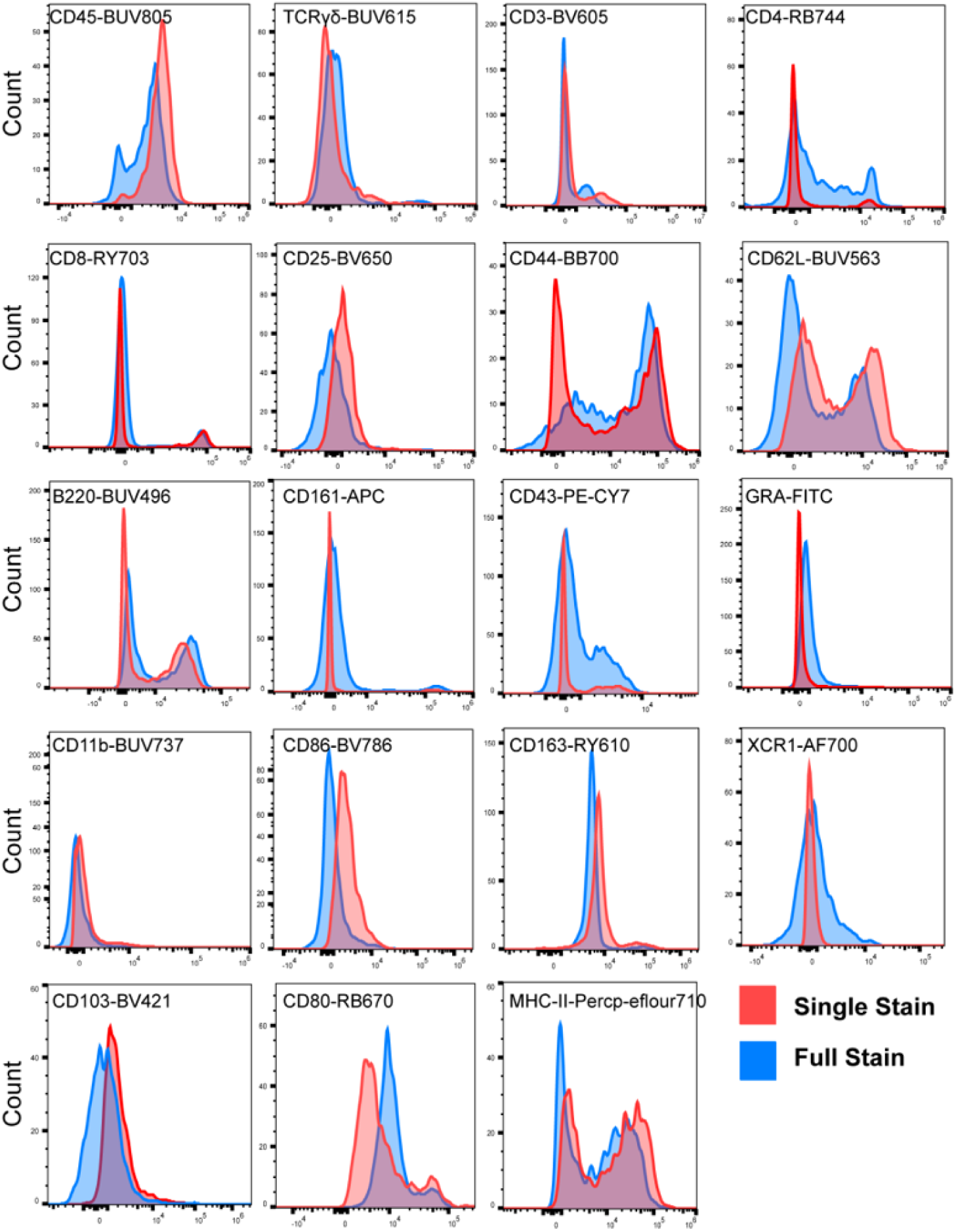
Comparison of fluorescence intensity between single-stained and fully stained samples for each marker after spectral unmixing. Single stained samples were marked with red line and full stained samples were marked with blue line. Through singlets, scatter gates, and backbone markers, data for lymphocytes, monocytes, or granulocytes were acquired.

## 5 Gating & Cell Phenotype Definition

The gating strategies applied to blood was shown in Figure 1A and the gating strategies applied to spleen and lymph nodes are shown in Figure 13 and Figure 14 respectively.

**Figure 13.**
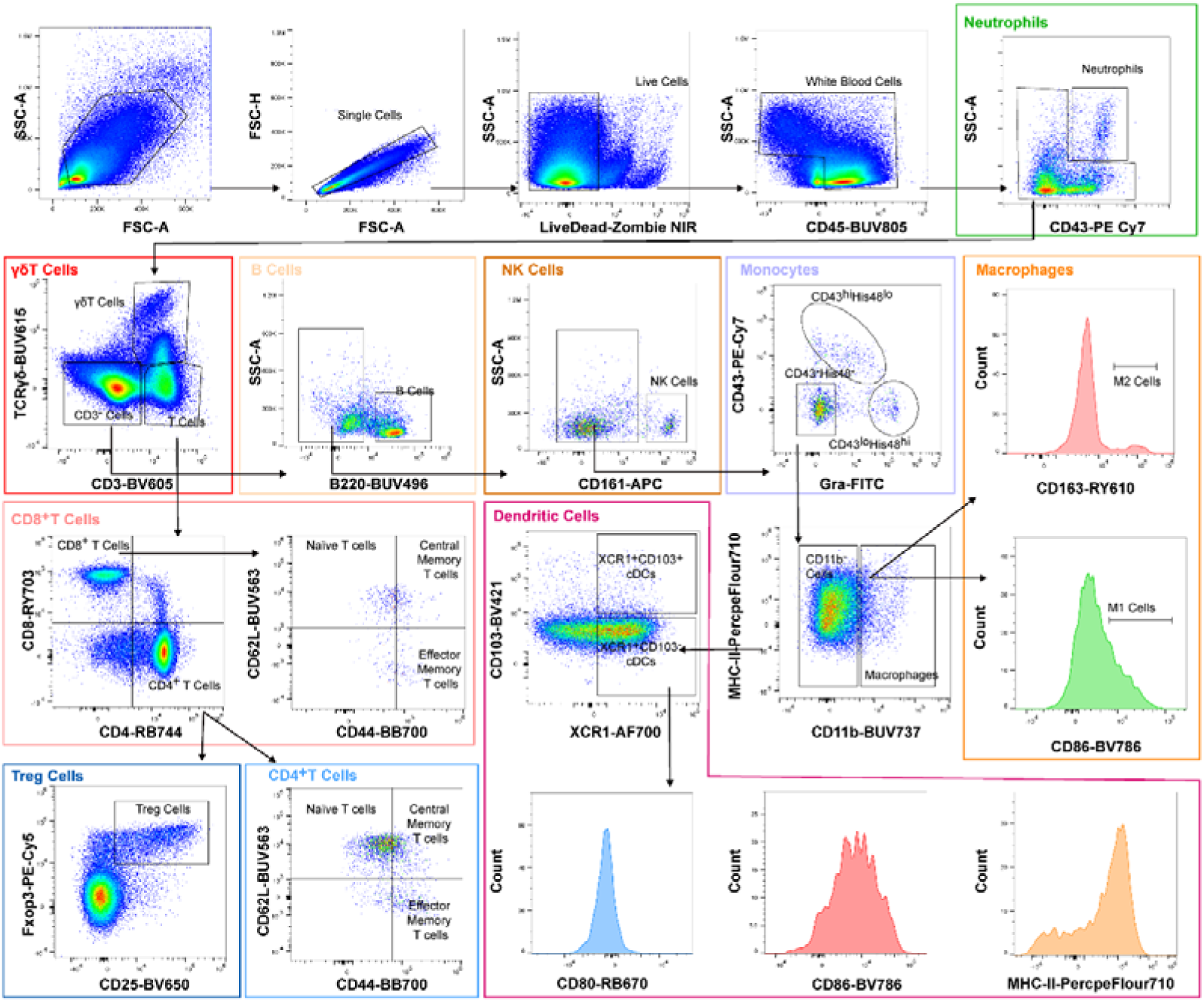
The 21-color gating strategy for the immune cells from spleen.

**Figure 14.**
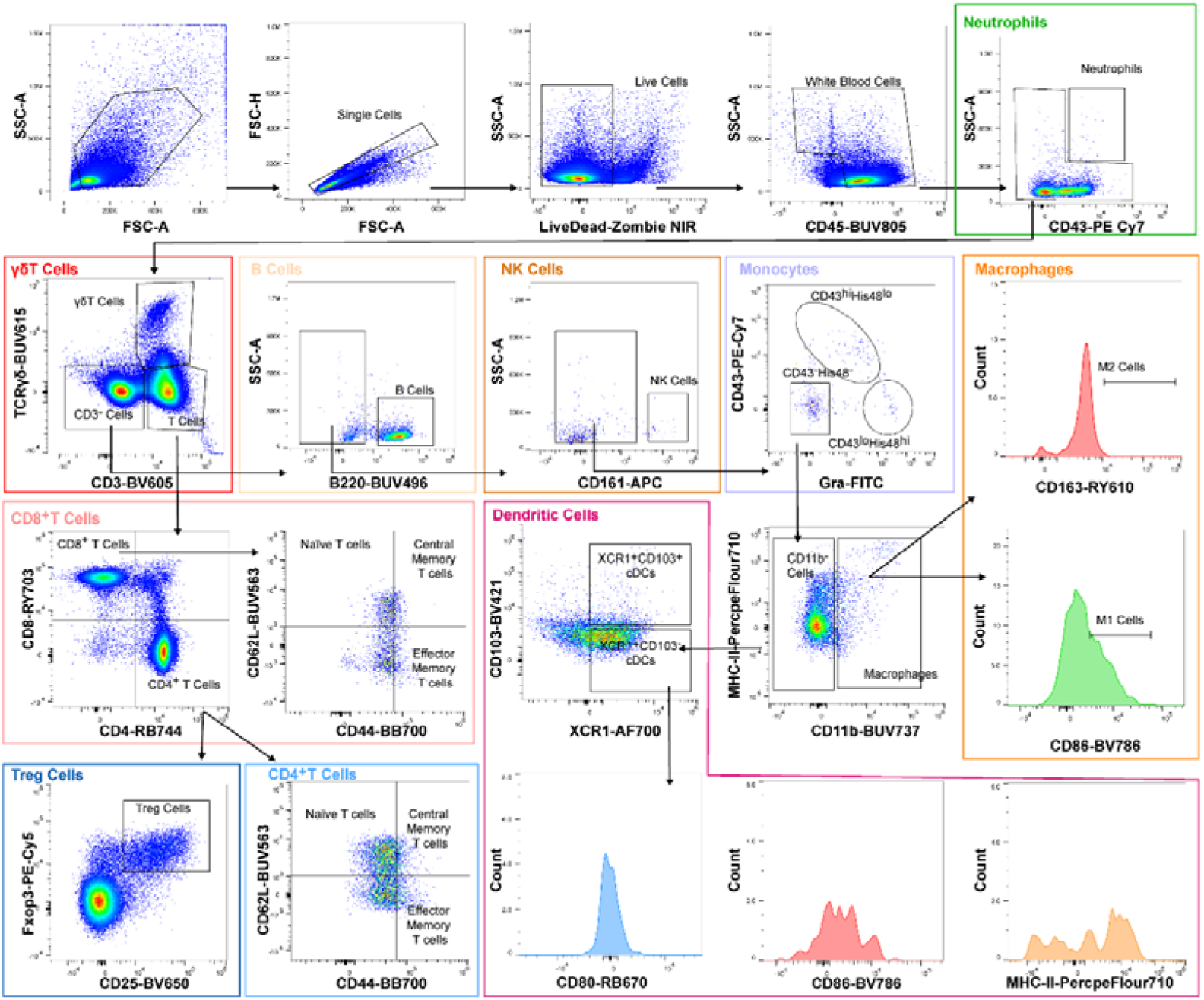
The 21-color gating strategy for the immune cells from lymph node.

## Author Contributions

Manuscript writing: Zhaoyuan Liang, Jie Zhang, Yuqing Wang

Experimental Design: Zhaoyuan Liang, Jie Zhang, Yuqing Wang

Data Analysis: Zhaoyuan Liang, Jie Zhang, Yuqing Wang, Yang Bai

Ethical Compliance: Xintong Zhang, Lixiang Xue

Manuscript Revision: Lixiang Xue, Jie Zhang,Yuqing Wang, Jianling Yang

Review and editing: All authors

Funding Acquisition: Lixiang Xue, Jie Zhang, Yuqing Wang

Supervision: Lixiang Xue

## Acknowledgments

This work was supported by National Clinical Research Center for Obstetrics and Gynecology of Peking University Third Hospital (BYSYSZKF2025005), Shanxi Key Laboratory of Digestive Diseases and Organ Transplantation (DDOT2025004), Peking University Third Hospital Clinical Key Project (BYSYZD2026089).

